# The Arabidopsis endosperm is a temperature-sensing tissue that implements seed thermoinhibition through phyB and PIF3

**DOI:** 10.1101/2022.06.13.495921

**Authors:** Urszula Piskurewicz, Maria Sentandreu, Gaëtan Glauser, Luis Lopez-Molina

## Abstract

Seed thermoinhibition, the repression of germination under high temperatures, prevents seedling establishment under potentially fatal conditions. Thermoinhibition is relevant for ecology, phenology and agriculture, particularly in a warming globe. The temperature sensing mechanisms and signaling pathways sustaining thermoinhibition are unknown. We found that thermoinhibition in *Arabidopsis thaliana* is not autonomously controlled by the embryo but is rather implemented by the endosperm surrounding the embryo. High temperature is sensed through endospermic phyB by accelerating its reversion from the active signaling Pfr form into the inactive Pr form, as described in seedlings. This leads to stabilization of endospermic PIF3, which represses the expression of the endospermic ABA catabolic gene *CYP707A1* and promotes endospermic ABA synthesis and release towards the embryo to block its growth. Furthermore, endospermic ABA represses embryonic PIF3 accumulation that would otherwise promote embryonic growth. Hence, under high temperatures PIF3 exerts opposite growth responses in the endosperm and embryo.

## Introduction

The onset of the embryo-to-seedling transition is a major event in the plant’s life cycle as the plant abandons its embryonic resistant state within the mature seed to form a fragile seedling. Subsequent reproductive success will depend on whether the seedling survives the environment it confronts. It is therefore unsurprising that germination control mechanisms have evolved to improve the seedling’s chances for survival.

Newly formed seeds exhibit primary seed dormancy (hereafter referred as dormancy), a trait whereby seeds do not germinate under otherwise favorable germination conditions ^1^. Over time, seeds lose dormancy as they age (a process referred as dry after-ripening), i.e. they acquire the capacity to germinate under favorable conditions, thus becoming non-dormant. Dormancy delays the onset of the embryo-to-seedling transition to ensure that seedlings are formed during a favorable season. However, non-dormant seeds continue to control their germination if confronted with non-favorable conditions. This includes to capacity to block seed germination under high temperatures, an adaptive trait referred as seed thermoinhibition. Seeds can be remarkably discerning when confronted with high temperatures and a difference of 1°C may determine whether a substantial percentage of a seed population germinates or not ^2^. Thermoinhibition furthers increases the odds of establishing a viable seedling since seedlings may not survive under persistent high temperatures. Hence, together with dormancy, thermoinhibition represents an additional control layer, enabling plants to form a seedling during a favorable season to conduct their reproductive phase. The trait is expected to have an impact on plant ecology, phenology and agriculture and this impact will be even greater as temperatures increase throughout the globe.

Thermoinhibition has been genetically studied in lettuce (*Lactuca sativa*) and *Arabidopsis thaliana* ^3,4, 5^. Thermoinhibition requires ABA synthesis, as mutants unable to synthesize ABA lack thermoinhibition, and is associated with low GA biosynthesis ^2, 3^. Toh et al. showed that *rgl2* mutants, lacking the GA response factor RGL2 have lower seed thermoinhibition although remaining responsive to high temperatures ^2^. Previous work showed that low GA levels lead to stabilization of DELLA factors, such as RGL2, which promote ABA synthesis ^6, 7^. Thus, other DELLA factors accumulating in *rgl2* mutants could still promote thermoinhibition by promoting ABA synthesis. However, and interestingly, Toh et al also reported that exogenous GA can promote germination at very high temperatures (34°C) but without fully abolishing thermoinhibition ^2^. This suggests that there are additional parallel pathways promoting thermoinhibition independently of the GA signaling pathway. Furthermore, the sensing mechanisms enabling seeds to trigger thermoinhibition is unknown.

Like seeds, seedlings are capable of discerning small changes in temperature to adapt their growth rate and avoid heat stress. Increasing temperatures promote hypocotyl and petiole growth as well as flowering ^8, 9^. Phytochromes are a family of five photoreceptors in Arabidopsis (phyA, phyB, phyC, phyD and phyE) synthesized in an signaling-inactive state known as Pr. Upon absorption of red light (R), they convert into a signaling-active Pfr state. In its Pfr state phyB interacts with phytochrome interacting factors (PIFs), a family of basic helix-loop-helix transcription factors regulating photomorphogenic development, to promote their degradation. In the Pr state PIFs are stabilized, which promotes their accumulation ^10, 11^. The Pfr state can revert to the Pr state upon absorption of far-red light (FR) or through temperature-dependent thermal relaxation (also called “thermal reversion”), a process accelerated by increasing temperatures and occurring even in absence of light ^10, 11^. Phytochrome thermal reversion, and particularly that of phytochrome phyB, was proposed to be the underlying mechanism by which seedlings sense temperature ^12, 13^. In turn, the signaling-inactive Pr state promotes the stabilization of phytochrome-interacting factors (PIFs) that promote growth ^10, 11^. Previous work also showed that phyB inactivation by far red-light in the seed endosperm blocks germination ^14^. Inactive phyB promotes ABA synthesis and release from the endosperm to block germination ^14, 15^. This conclusion was notably reached using a “Seed Coat Bedding Assay” (SCBA) whereby dissected embryos are cultured on a bed of dissected endosperms, which enables to study embryonic growth under the influence of endosperm tissues ^16^. The SCBA enables the use of embryo and endosperm of different genetic backgrounds to genetically dissect the processes regulating growth in the embryo and endosperm. However, whether phytochromes and the endosperm play a role in seed thermoinhibition is poorly understood.

Here, we show that thermoinhibition in *Arabidopsis thaliana* is entirely dependent on the endosperm rather than the embryo as embryos deprived of their endosperm are unable to repress their growth under high temperatures. Using a SCBA and direct ABA measurements, we show that at high temperatures the endosperm releases ABA towards the embryo to block its growth and maintain its mature state by promoting the accumulation of the ABA response factor ABI5. In the endosperm, high temperature reduces the pool of active phyB, which activates endospermic ABA synthesis and release. This involves endospermic DELLA factors and PIFs, and particularly PIF3, which represses the expression of *CYP707A1*, encoding an ABA catabolic gene (Kushiro et al. 2004). We found that PIF3 strongly accumulates in the embryo during the embryo-to-seedling transition and promotes early seedling growth. Furthermore, under high temperatures endospermic ABA represses embryonic PIF3 accumulation, indicating that PIF3 exerts opposite growth responses in the endosperm and embryo under high temperatures.

## Results

### The endosperm enables seed thermoinhibition by releasing ABA towards the embryo

The endosperm promotes dormancy by releasing ABA towards the embryo upon imbibition. We explored whether the endosperm plays a role for seed thermoinhibition. Embryos deprived of their seed coat and endosperm 4 hours after seed imbibition grew under high temperatures (34°C), expanding their cotyledons, doubling their hypocotyl length 3 days after dissection and eventually dying (Fig. 1a and b). In contrast, embryos from intact seeds did not germinate and maintained their embryonic state (Fig. 1a and b). Furthermore, embryos remained thermoinhibited when only the seed coat was removed (Fig. 1c). These results show that presence of the endosperm is essential for seed thermoinhibition. Previous work showed that endogenous ABA synthesis is necessary for thermoinhibition but whether the ABA synthesis in the endosperm contributes to thermoinhibition is not known.

**Figure 1.**
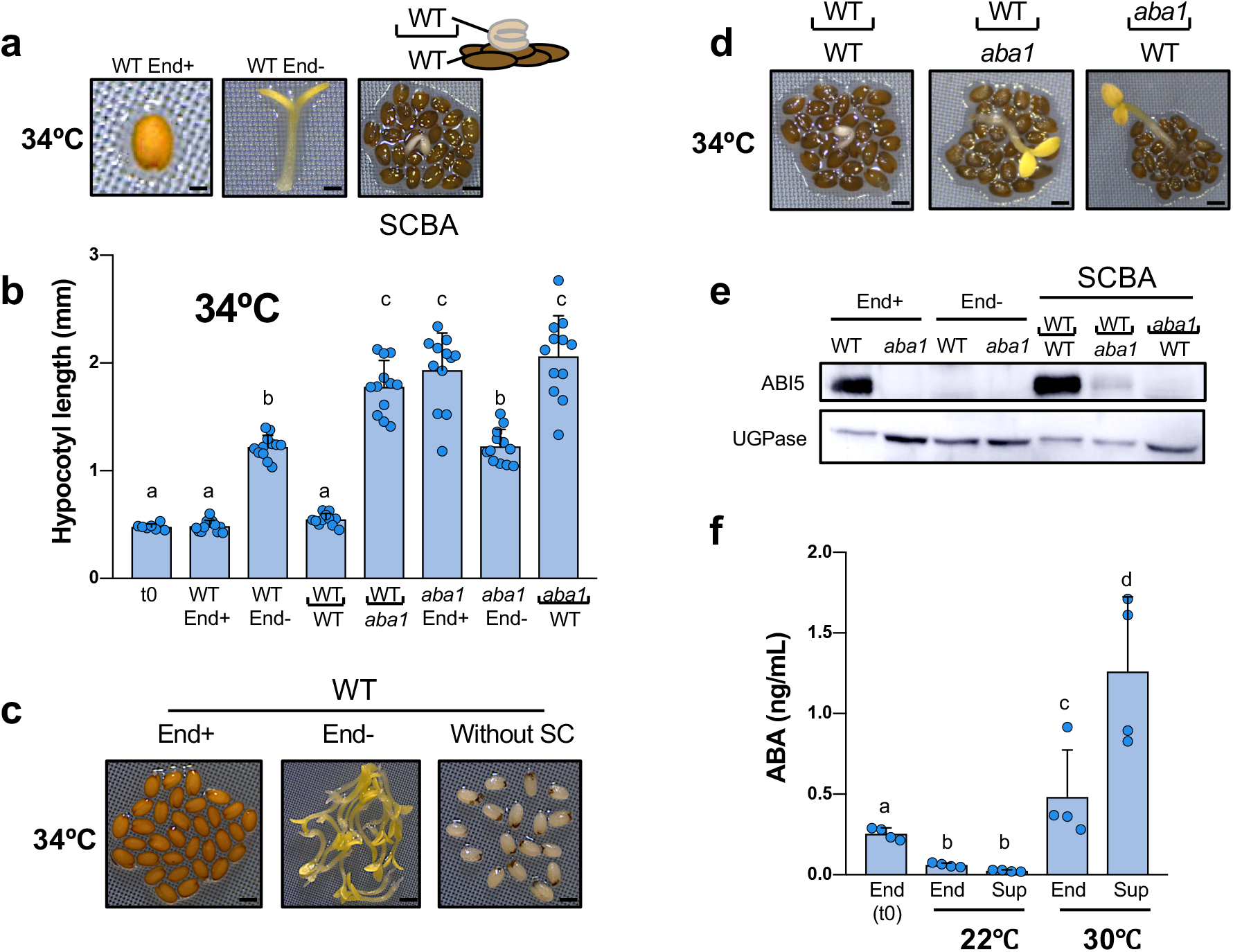
The endosperm enables seed thermoinhibition by releasing ABA towards the embryo. a. Representative pictures taken 3 days after seed imbibition at 34°C of a WT seed (End+), a WT embryo upon removal of the endosperm 4 hours after seed imbibition (End-) and a WT embryo cultured on a bed of WT endosperms with the seed coat still attached (seed coat bedding assay -SCBA-)(bar, 0.1mm, 0.3 mm and 0.5 mm for left, center and right picture, respectively). b. Hypocotyl length (mm) of WT and *aba1* embryos dissected 4 hours after seed imbibition and cultured for 3 days at 34°C without (End+) or with endosperm (End-) dissection. For the SCBAs, WT or *aba1* embryos were cultured on a bed 30 WT or *aba1* endosperms as indicated (n=15). Lower case letters above histograms are used to establish whether two values are statistically significantly different as assessed by one-way ANOVA followed by a Tukey HSD test (p<0.05): different letters denote statistically different values. c. Representative pictures taken 3 days after seed imbibition at 34°C of WT seeds (End+; bar: 0.5 mm), WT embryos upon removal of the endosperm (End-; bar: 1 mm) or of only the seed coat (Without SC; bar: 0.5 mm) 4 hours after seed imbibition. d. Representative pictures taken 3 days after seed imbibition at 34°C of WT and *aba1* embryos dissected 4 hours after seeds imbibition and cultivated on a bed of WT or *aba1* endosperms as indicated. Bar: 0.5 mm (left, center) and 0.8 mm (right). Hypocotyl length quantification is shown in b. e. ABI5 protein levels in WT and *aba1* embryos dissected from seeds (End+) 3 days upon seed imbibition and in WT and *aba1* embryos dissected 4h after seed imbibition and further cultured for 3 days (End-) at 34°C. For the SCBAs, WT or *aba1* embryos were cultured on a bed of WT or *aba1* endosperms as indicated. UGPase protein levels were used as a loading control. f. WT endosperms with the seed coat still attached were dissected 4 hours after seed imbibition and cultured in water at 22 or 30°C for 40 hours. Histograms show ABA levels measured in the endosperm tissue at the time of dissection (t0), 40 hours after incubation (End) and in culture medium (Sup). Data from 4 independent experiments (500 endosperms per experiment). Statistical differences assessed by Student’s t-test (p<0.05): different letters denote statistically different values.

In a seed coat bedding assay (SCBA), embryos are cultured on a bed of dissected endosperms (with the seed coat still attached) to study the influence of the endosperm on embryonic growth ^16^. WT embryos cultured on a bed of WT endosperms under high temperatures were unable to grow, maintaining the same appearance as embryos within intact seeds, and accumulated high levels of the ABA response transcription factor ABI5 (Fig. 1a, b, d and e) ^17^. Furthermore, WT embryos cultured on a bed of *aba1* endosperms, unable to synthesize ABA, were able to grow and had low ABI5 accumulation (Fig. 1b,d and e). These observations strongly indicated that seed thermoinhibition involves ABA release by the endosperm towards the embryo to block its growth.

Consistent with this view, endosperms cultured at 30°C for 48h contained 8-fold more ABA than those cultured at 22°C (Fig. 1f). Furthermore, over the same period, the levels of ABA in the culture medium at 30°C were more than 16x higher than those at 22°C (Fig. 1f). These observations show that cultured dissected endosperms actively synthesize and release ABA at 30°C but not at 22°C. Interestingly, *aba1* mutant embryos were able to grow when cultured on a bed of WT endosperms and did not accumulate ABI5 (Fig. 1b, d and e). Altogether, these results support the notion that 1) the endosperm is necessary to implement thermoinhibition by synthesizing and releasing ABA towards the embryo and 2) that ABA synthesized in the embryo also participates to promote thermoinhibition without being sufficient.

### High temperature promotes thermoinhibition by repressing phyB signaling in the endosperm

Previous work showed that the endosperm blocks the germination of non-dormant seeds in response to FR light. Indeed, in the endosperm FR light converts the phytochrome B (phyB) Pfr signaling active form into its Pr inactive form, which promotes synthesis of ABA and its release towards the embryo to block its germination ^14^. Phytochrome signaling in seedlings was shown to be affected by temperature. Indeed, increasing temperatures promotes Pfr reversion into the Pr inactive form and promotes hypocotyl growth ^12, 13^. We therefore hypothesized that high temperatures block germination by repressing phyB signaling in the endosperm.

In the Col-0 accession, seed thermoinhibition in six months-old seeds is weak at 28°C and is better observed at temperatures in the 30-34°C range (Fig. 2a)(see discussion). At 34°C, seed germination was blocked in WT, *phyB* and *phyAphyCphyDphyE* (*phyACDE*) mutant seeds (Fig. 2a). In contrast, at 28°C WT and *phyACDE* seeds germinated whereas *phyB* mutants did not germinate, consistent with a previous report (Fig. 2a) ^18^. At 22°C, *phyB* and *phyACDE* mutant seeds germinated with the same percentage as WT seeds. Hence, seeds deficient in phyB signaling maintain seed thermoinhibition when temperature drops from 34°C to 28°C, unlike WT seeds. This suggests that absence of seed thermoinhibition at 28°C in WT seeds requires active phyB signaling and, conversely, that thermoinhibition at 34°C in WT seeds could be due to inactive phyB signaling. At 22°C, *phyB* mutants germinated, suggesting that other phytochromes promote germination at lower temperatures (see discussion).

**Figure 2.**
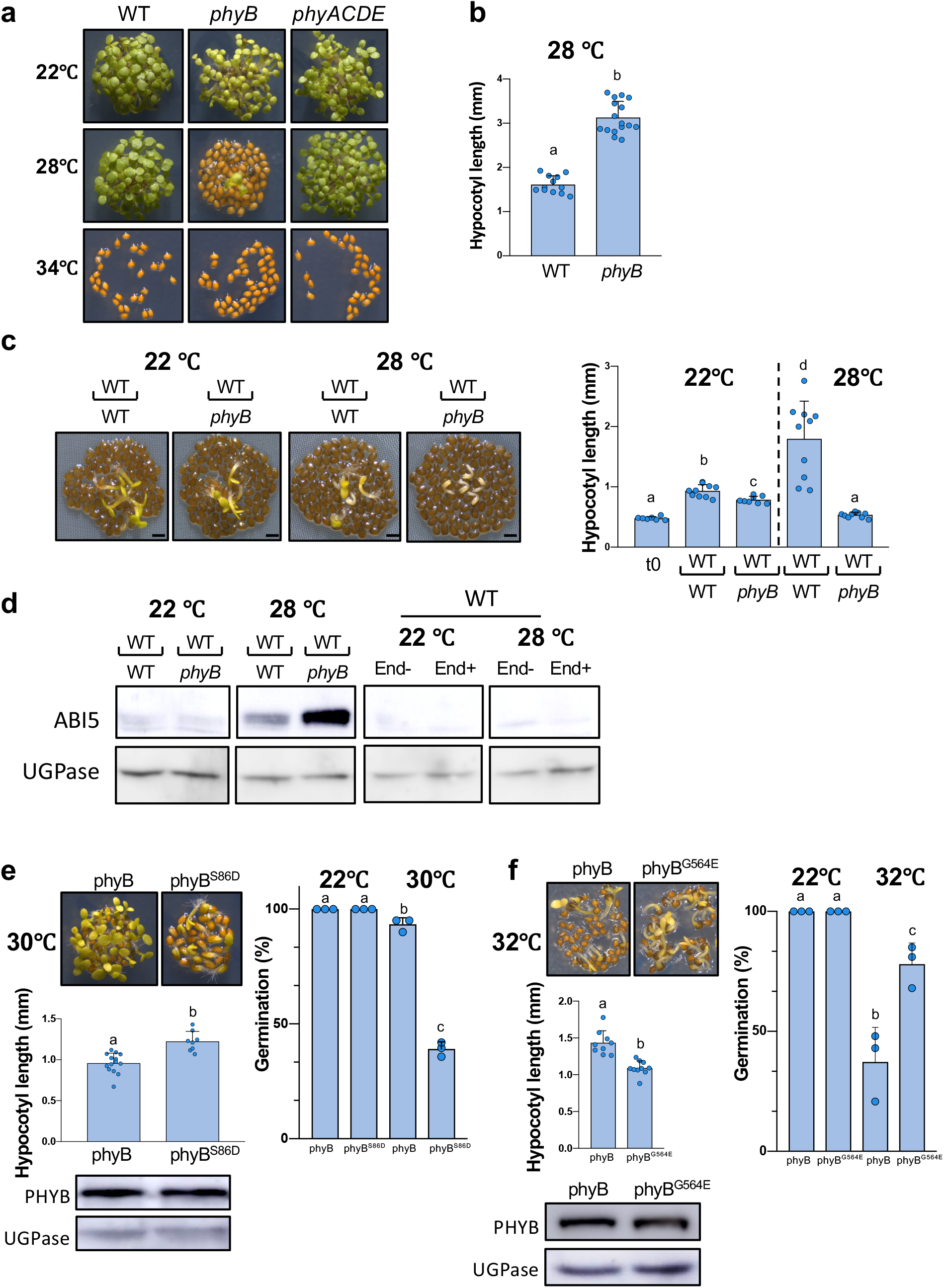
High temperatures promote thermoinhibition by repressing phyB signaling in the endosperm. a. Representative pictures taken 4 days after seed imbibition of WT, *phyB* and *phyACDE* seeds cultivated at 22°C, 28°C or 34°C. The experiment was repeated three times giving the same results. b. Hypocotyl length (mm) of WT and *phyB* embryos dissected 4 hours after seed imbibition and cultured for 4 days at 28°C (n=12 embryos). Statistical treatment and lower-case letters as in Figure 1b. c. Left panel: representative pictures taken 3 days after seed imbibition at 22°C and 28°C of WT embryos dissected 4 hours after seed imbibition and cultivated on a bed of WT or *phyB* endosperms as indicated. Bar: 0.8 mm. Right panel: hypocotyl length (mm) after 3 days (n=10 embryos). Statistical treatment and lower-case letters as in Figure 1b. d. First two panels: ABI5 protein levels 3 days upon seed imbibition at 22°C or 28°C in WT embryos cultured on a bed of WT or *phyB* endosperms as indicated. Last two panels: ABI5 levels in embryos 3 days upon seed imbibition at 22°C or 28°C in WT seeds (End+) and embryos upon removal of the endosperm 4h after seed imbibition (End-). UGPase protein levels were used as a loading control. e. Representative pictures taken 3 days after seed imbibition at 30°C of *phyB* mutant seeds complemented with phyB that has a WT Ser-86 (phyB) or Ser86Asp (phyB^S86D^). Average germination percentage after 3 days at 22°C and 30°C in 3 independent seed batches (n≥50 seeds). Statistical differences assessed by Student’s t-test. Hypocotyl length (mm) of embryos dissected 4 hours after seed imbibition and cultured for 3 days at 30°C (n≥8 embryos). Statistical treatment and lower-case letters as in Figure 1b. Protein gel blot analysis of phyB protein levels in seeds of the two phyB lines incubated for 24 hours at 30°C. UGPase protein levels were used as a loading control. f. Same as 2e using *phyB* mutant seeds complemented with phyB that has a WT Gly564 (phyB) or Gly564Glu (phyB^G564E^).

Strikingly, at 28°C *phyB* embryos deprived of their endosperm were able to grow, even reaching a longer hypocotyl length relative to WT embryos after 4 days, despite the strong thermoinhibition observed in intact *phyB* mutant seeds (Fig. 2b). The longer hypocotyl of *phyB* mutant seedlings arising from endosperm-less embryos is consistent with previous work showing that phyB signaling represses hypocotyl elongation in seedlings ^11^. Furthermore, WT embryos cultured on a bed of *phyB* mutant endosperm at 28°C had their growth strongly repressed in comparison to that of WT embryos cultured on a bed of WT endosperms and accumulated higher ABI5 protein levels (Fig. 2c and d). At 22°C, WT and *phyB* endosperms were unable to block the growth of WT embryos nor to trigger high ABI5 accumulation (Fig. 2c and d). Hence, these observations strongly indicate that seed thermoinhibition of *phyB* mutant seeds at 28°C is implemented by higher endospermic ABA release relative to WT endosperm as a result of absent phyB signaling.

Previous reports have shown that phyB thermal reversion can be accelerated or slowed down by specific amino acid changes. We compared seed thermoinhibition in a *phyB* mutant complementation line in which thermal reversion is accelerated by a Ser86Asp amino acid change (phyB^S86D^) relative to a control line bearing the WT phyB sequence (phyB)(materials and methods) ^19^. At 30°C, the phyB^S86D^ line had higher thermoinhibition than phyB even though both lines accumulated similar amounts of transgenic gene products (Figure 2e). Stronger thermoinhibition in the phyB^S86D^ line is most likely the result of stronger endosperm-imposed germination arrest because the growth of phyB^S86D^ embryos was faster than that of phyB embryos upon endosperm removal (Figure 2e). We also assessed thermoinhibition in *phyB* mutant transgenic lines in which phyB thermal reversion is slowed down by a G564E amino acid change (phyB^G564E^) relative to a control line bearing no change (phyB) ^20, 21^. At 32ºC the phyB^G564E^ line was less thermoinhibited than the phyB control line even though the hypocotyl of the control line elongated faster than that of the phyB^G564E^ line (Figure 2f). The complemented lines accumulated similar amounts of transgenic gene products at 32ºC (Figure 2f). Altogether, these results are consistent with a seed thermoinhibition model whereby high temperatures promote endospermic phyB thermal reversion, which leads to higher endospermic ABA synthesis and release towards the embryo to block its growth.

### DELLA factors promote endospermic ABA synthesis and release when high temperatures reduce phyB signaling

We next sought to identify the endospermic factors promoting ABA synthesis and release when phyB signaling is repressed in response to high temperatures. The DELLA factors were potential candidates for several reasons: 1) low phyB signaling represses the expression of GA biosynthesis genes, 2) the DELLA factors RGL2, GAI and RGA promote ABA synthesis when GA levels are low and 3) *rgl2* mutants seeds have low seed thermoinhibition responses ^2, 6, 7, 22^. We first evaluated whether endospermic DELLA factors promote thermoinhibition. At 32°C, *della* mutants seeds, bearing loss-of-function mutations in all five *Arabidopsis* DELLA genes (*RGL2, GAI, RGA, RGL1* and *RGL3*), lacked seed thermoinhibition unlike WT seeds (Supplementary Figure 1a). Furthermore, at 32°C, *della* endosperms were unable to repress WT embryo growth nor induce strong embryonic ABI5 accumulation in a SCBA unlike WT endosperms (Figure 3a). Altogether, these observations strongly suggest that endospermic DELLA factors promote thermoinhibition by enhancing endospermic ABA synthesis and release under high temperatures.

**Figure 3.**
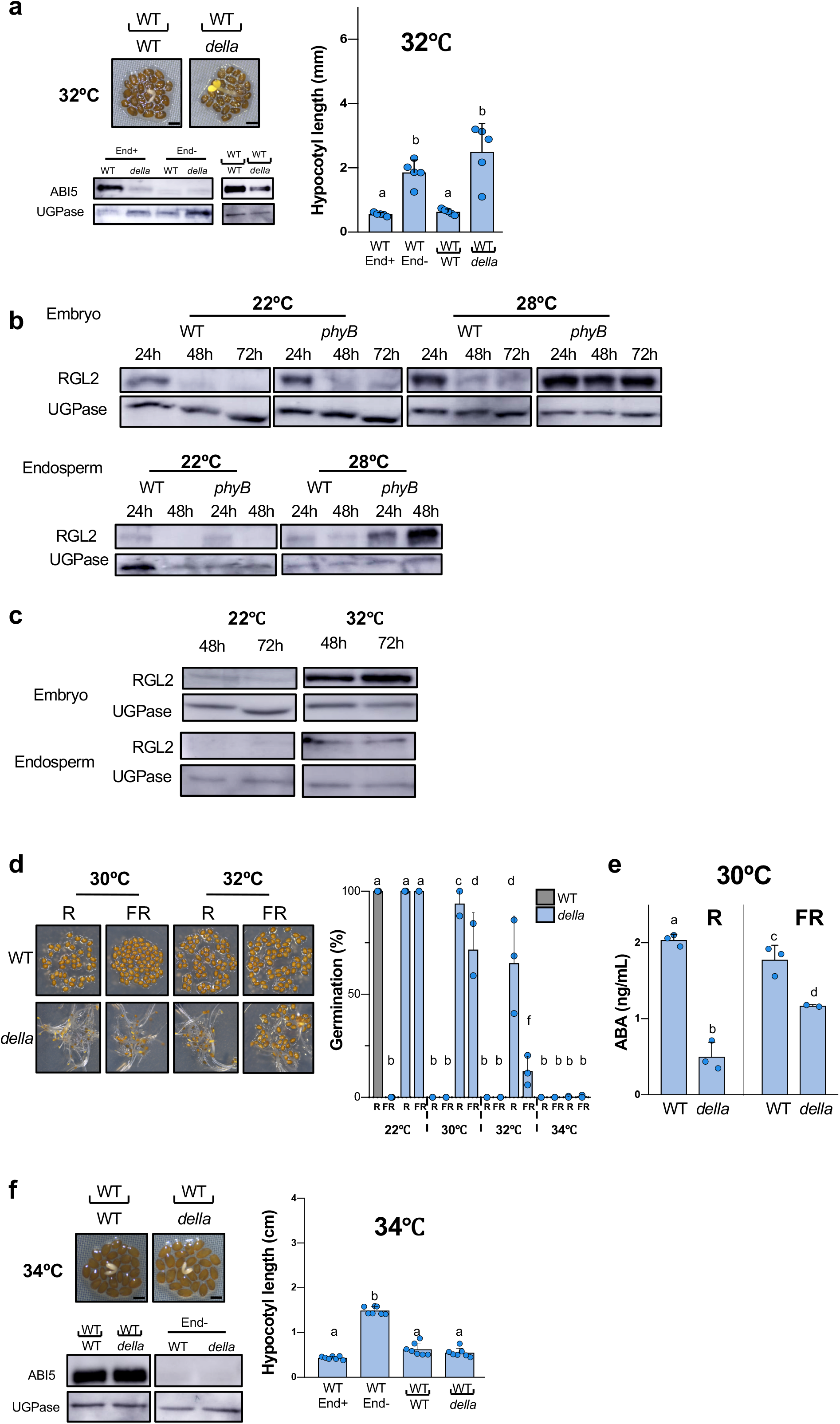
DELLA factors promote endospermic ABA synthesis and release when high temperatures reduce phyB signaling. a. Left panel: representative pictures taken 3 days after seed imbibition at 32°C of WT embryos dissected 4 hours after seeds imbibition and cultivated on a bed of WT or *della* endosperms as indicated. Bar: 0.5 mm. Right panel: hypocotyl length (mm) of WT embryos dissected 4 hours after seed imbibition and cultured for 3 days at 32°C without (End+) or with (End-) endosperm dissection. For the SCBAs, WT embryos were cultured on a bed of WT or *della* endosperms as indicated (n=5). Bottom panel: ABI5 protein levels in WT and *della* embryos dissected from seeds (End+) 3 days upon seed imbibition and in WT and *della* embryos dissected 4h after seed imbibition and further cultured for 3 days (End-) at 32°C. For the SCBAs, WT embryos were cultured on a bed of WT or *della* endosperms as indicated. Each lane contained protein extracts from 10 embryos. UGPase protein levels were used as a loading control. b. Protein gel blot analysis of RGL2 levels in WT and *phyB* embryos and endosperms cultivated at 22 and 28°C and dissected at the indicated times upon seed imbibition. UGPase protein levels were used as a loading control. c. Same as 3b using WT seeds cultivated at 22°C and 32°C. d. Representative pictures taken 3 days after seed imbibition of WT and *della* seeds cultivated in darkness at 30°C or 32°C after receiving a far-red light pulse (FR) or a FR pulse followed by a red light pulse (R) 2h upon seed imbibition. Average germination percentages after 3 days at 22°C, 30°C, 32°C and 34°C in 3 independent seed batches (n≥50 seeds). Statistical differences assessed by Student’s t-test. e. WT and *della* endosperms were dissected 4 hours after seed imbibition from WT and *della* seeds irradiated with FR/R or FR 2h upon seed imbibition and cultured in water in darkness at 30°C for 40 hours. Histograms show ABA levels measured in culture medium after 40 hours. Data from 3 independent experiments (150 endosperms per experiment). Statistical differences assessed by Student’s t-test. f. Same as Figure 3a but seeds were incubated at 34°C.

We next evaluated the hypothesis that DELLA factors promote endospermic ABA when high temperatures reduce endospermic phyB signaling. phyB signaling promotes synthesis of GA, which binds to the GID1 receptors leading to their interaction with DELLA factors and promotion of DELLA proteasomal degradation ^23, 24, 25, 26^. At 22°C, RGL2 protein levels were similar in the endosperm and embryo of WT and *phyB* seeds (Figure 3b). At 28°C, a temperature that blocks the germination of *phyB* seeds but not that of WT seeds, RGL2 levels marginally increased in the endosperm and embryo of WT seeds (Figure 3b). In contrast, at 28°C RGL2 levels strongly increased in both tissues of *phyB* mutant seeds (Figure 3b). At 32°C, a temperature blocking WT seed germination, RGL2 levels strongly increased in the endosperm and embryo of WT seeds (Figure 3c).

phyB accumulation was similar in WT and *della* mutant seeds indicating that enhanced phyB signaling in *della* seeds does not account for their lack of thermoinhibition (Supplementary Figure 1). *della* mutant seeds irradiated with FR light upon imbibition, which inactivates phyB, and maintained at 30°C in darkness continued to lack seed thermoinhibition, unlike WT seeds (Figure 3d). Consistent with this result, at 30°C, FR-irradiated *della* endosperm released 1.5-fold lower ABA levels relative to WT endosperm (Figure 3e). In addition, under white light *della/phyB* mutant seeds, lacking DELLA factors and phyB, were not thermoinhibited at 28°C, unlike *phyB* mutant seeds (Supplementary Figure 1c). Altogether, these observations support the notion that DELLA factors promote thermoinhibition when phyB signaling is repressed by high temperatures.

### phyB inactivation by high temperatures promotes seed thermoinhibition even in absence of DELLA factors

Interestingly, however, several observations suggested that upon lowering of phyB signaling by high temperatures another endospermic pathway promotes ABA synthesis and release independently of DELLA factors. Indeed, 1) thermoinhibition could be observed at 30°C in about a quarter of *della* seeds irradiated with FR, which inactivates phyB and, furthermore, the percentage of thermoinhibited *della* seeds further increased with increasing temperatures, which accelerates phyB reversion, reaching 100% at 34°C (Figure 3d); 2) *dellaphyB* mutant seeds, lacking DELLA factors and phyB, were more thermoinhibited at 30°C than *della* mutant seeds (Supplementary Figure 1d); 3) At 30°C, FR-irradiated *della* endosperms continued to release substantial amounts ABA even though they were lower than those released by FR-irradiated WT endosperms (Figure 3e); 4) at 34°C in presence of white light, a substantial percentage (40%) of *della* seeds were thermoinhibited whereas no thermoinhibition took place in *aba1* seeds, deficient in ABA synthesis (Supplementary Figure 1a) and 5) *della* seed thermoinhibition required the presence of the endosperm and, accordingly, at 34°C, *della* endosperms were able to repress the growth of WT embryos and induce ABI5 accumulation in a SCBA (Figure 3f). Finally, Toh et al observed that WT seed thermoinhibition at 34°C was not fully abolished in presence of exogenous GA, which promotes DELLA factor degradation ^2^. We therefore sought to identify additional endospermic factors promoting ABA synthesis and release.

### Endospermic PIF3 promotes seed thermoinhibition by promoting ABA synthesis and release

In its active Pfr form phyB interacts with PIFs transcription factors to promote their degradation so that when phyB is in its inactive Pr form PIFs are stabilized, which promotes their accumulation ^10, 11^. We therefore asked whether PIFs could promote seed thermoinhibition. *pifq* mutant seeds, lacking PIF1, PIF3, PIF4, PIF5, PIF7, were thermoinhibited at 32°C, indicating that PIFs are not essential for seed thermoinhibition at very high temperatures (Supplementary Figure 1e). We therefore monitored seed thermoinhibition responses at 30°C, a moderately high temperature that does not fully promote thermoinhibition in WT seeds (Fig. 4a). At 22°C, no noticeable differences in seed germination were observed between WT and various single or multiple *pif* mutant seeds (Supplementary Fig. 2a). At 30°C, *pif4, pif6* and *pif7* seed thermoinhibition was similar to that of WT seeds. *pif1* and *pif5* seed thermoinhibition was lower than WT in some seed batches (Batches 1 and 2) but not in others (Batches 3 and 4)(Fig. 4a). However, *pif3* thermoinhibition was consistently lower in all seed batches, suggesting that PIF3 plays a significant role to promote thermoinhibition among other PIFs. Combining the *pif3* mutation with *pif1* and/or *pif5* mutations lead to even lower thermoinhibition in all seed batches (Fig. 4a). The double mutant *pif3pif4* was also less thermoinhibited than *pif3* suggesting that PIF4 could also promote thermoinhibition in absence of PIF3. At higher temperatures (32°C), these same *pif* mutant seeds increased their thermoinhibition, suggesting that DELLA factors overcome the absence of PIF1, PIF3 and PIF5 to promote thermoinhibition (Supplementary Fig. 2b). Consistent with this hypothesis, *pif1, pif5* and *pif3* mutant thermoinhibition in presence of GA (5 µM) strongly decreased relative to WT at 32°C and even further decreased in the double and triple mutant combinations (Supplementary Fig. 2b). Altogether, these data show that PIFs, and notably PIF3, promote seed thermoinhibition. Together with the observation that *pifq* mutants are thermoinhibited at very high temperatures (Supplementary Figure 1e), our results indicate that DELLA factors and PIFs promote endospermic ABA synthesis and release through parallel pathways. Hereafter, we focus on the role of PIF3 to promote thermoinhibition. PIFs, including PIF3, were previously shown to promote hypocotyl elongation, which contrasts with their activity to promote seed thermoinhibition, which blocks the embryo-to-seedling transition ^10, 11, 27^. This is consistent with the notion that PIF3 thermoinhibition activity takes place in the endosperm, rather than in the embryo, by promoting ABA synthesis and release.

**Figure 4.**
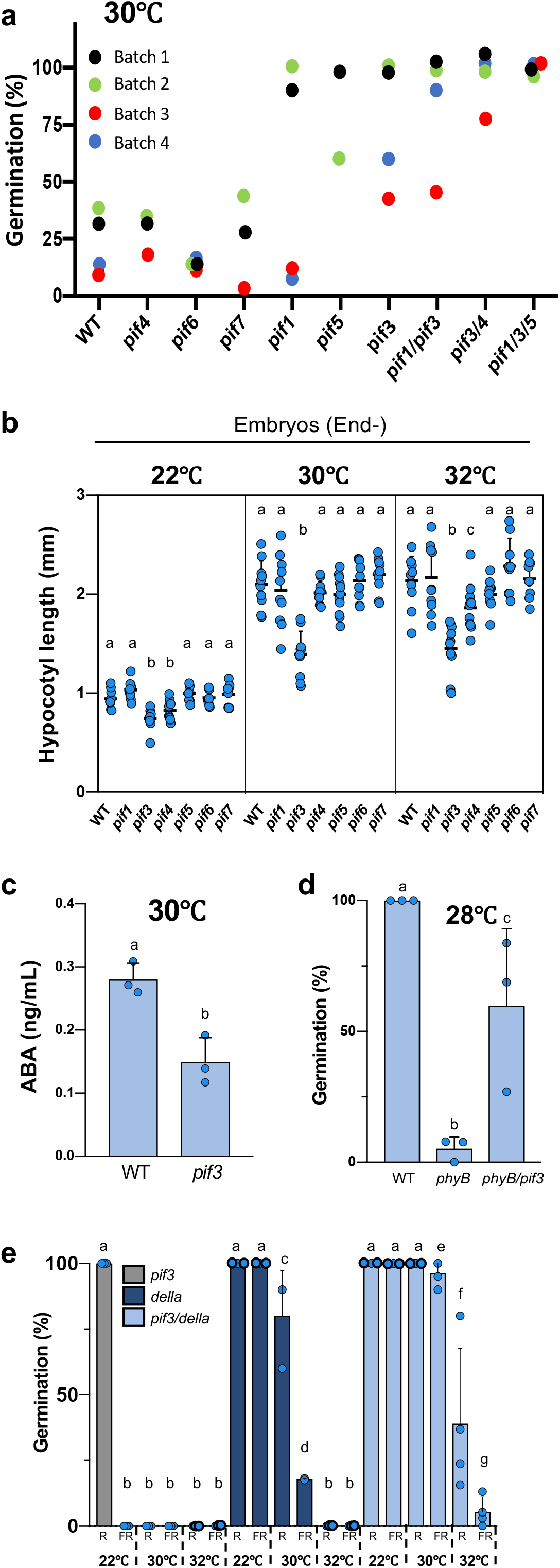
Endospermic PIF3 promotes seed thermoinhibition by promoting ABA synthesis and release. a. Average germination percentages of WT seeds and seeds carrying mutations in *PIF* genes, as indicated, after 4 days at 30°C in 4 independent seed batches (n≥50 seeds). b. Hypocotyl length (mm) of WT and *pif* embryos dissected 4 hours after seed imbibition and cultured for 3 days at 22°C, 30°C and 32°C (n=10). Statistical treatment and lower-case letters as in Figure 1b. c. WT and *pif3* endosperms were dissected 4 hours after seed imbibition from WT and *pif3* seeds and cultured in water at 30°C for 40 hours. Histograms show ABA levels measured in culture medium after 40 hours. Data from 3 independent experiments (150 endosperms per experiment). Statistical differences assessed by Student’s t-test. d. Average germination percentages of WT, *phyB* and *phyB/pif3* seeds after 3 days at 28°C in 3 independent seed batches (n≥50 seeds). Statistical differences assessed by Student’s t-test. e. Average germination percentages of *pif3, della* and *pif3/della* seeds cultivated in darkness for 3 days at 22°C, 30°C and 32°C after receiving a far-red light pulse (FR) or a FR pulse followed by a red light pulse (R) 2h upon seed imbibition in 3 independent seed batches (n≥50 seeds). Statistical differences assessed by Student’s t-test.

To verify these claims, we sought to 1) assess embryonic PIF3 activity to promote early seedling growth in embryos deprived of their endosperm and 2) assess endospermic PIF3 activity to promote ABA synthesis and release.

We measured the hypocotyl length of 4-day-old WT and different *pif* mutant seedlings derived from embryos whose endosperm was removed 4h upon seed imbibition (Fig. 4b). Hypocotyls elongated faster at 30°C and 32°C than at 22°C in all genotypes tested, consistent with previous reports (Fig. 4b)^8, 28^. Interestingly, relative to WT, the hypocotyl of the *pif3* mutant was markedly shorter at 30°C (34% shorter) and 32°C (32% shorter) and to a lesser extent at 22°C (20% shorter) whereas that of the other *pif* mutants was similar with the exception of *pif4*, which was slightly shorter at 22°C (12% shorter) and 32°C (14% shorter)(Fig. 4b). Koini et al. examined hypocotyl elongation of 4-day-old seedlings upon transfer to high temperatures (28°C) and found that the hypocotyl of *pif4* seedlings elongated less than WT, whereas that of *pif3* seedlings responded normally ^8^. We could confirm these observations and, together with our observations, we conclude that PIF3 plays a specific role to promote early hypocotyl elongation particularly at high temperatures (Supplementary Fig. 2c).

Next, we assessed PIF3 activity in the endosperm to promote ABA synthesis and release. We dissected WT and *pif3* endosperms 4 hours after seed imbibition and cultured them in water at 30°C for 48 hours. The *pif3* culture medium contained significantly lower ABA levels than that of the WT culture medium (Fig. 4c). This result shows that PIF3 activity in endosperm is necessary to release ABA at high temperatures. Consistent with this conclusion, *pif3/phyB* seeds were less thermoinhibited than *phyB* seeds at 28°C (Fig. 4d). Moreover, the *della/pif3* mutant was less thermoinhibited at 30ºC and 32ºC than the *della* quintuple mutant showing that PIF3 can promote thermoinhibition independently of DELLA factors (Figure 4e)(materials and methods). Altogether, these results corroborate the view that PIF3 promotes endospermic ABA synthesis and release when phyB signaling is repressed by high temperatures. Furthermore, these findings also lead us to conclude that in absence of the endosperm, PIF3 indeed promotes early seedling growth under high temperatures. This conclusion is rather intriguing since on one hand, PIF3 promotes embryonic growth in absence of the endosperm but on the other it represses growth in presence of the endosperm. To better understand this apparent paradox, we sought to understand how PIF3 protein accumulation is regulated under high temperatures in the endosperm and embryo.

### Endospermic ABA represses embryonic *PIF3* expression

Remarkably, in WT seeds imbibed at 22°C in presence of white light PIF3 protein accumulation increased markedly but transiently at 36h and 48h in the embryo upon seed imbibition, which is the time when the embryo is germinating, and the hypocotyl of the seedling being formed rapidly elongates (Fig. 5a). This is consistent with the role of PIF3 to promote early seedling growth as shown above. In contrast, at 34°C, PIF3 embryonic accumulation remained low over time upon imbibition (Fig. 5a). However, endospermic PIF3 accumulation was lower at 22°C that at 34°C (Fig. 5a). A transgenic line carrying a *PIF3* promoter sequence fused to GUS (prom*PIF3::GUS*) yielded results consistent with those of PIF3 accumulation: at 22°C, embryonic GUS activity increased strongly between 24h and 48h upon imbibition and diminished thereafter whereas it was undetected in thermoinhibited embryos at 34°C (Fig. 5b). Conversely, endosperm GUS activity was higher at 34°C than at 22°C (Fig. 5b). This suggests that at 22°C the observed increases in embryonic PIF3 accumulation are driven by increased embryonic *PIF3* mRNA accumulation whereas at 34°C, low embryonic PIF3 levels are due to low embryonic *PIF3* mRNA levels. In contrast to the embryo, *PIF3* mRNA accumulation in the endosperm is stimulated by high temperatures, which promotes endospermic PIF3 accumulation.

**Figure 5.**
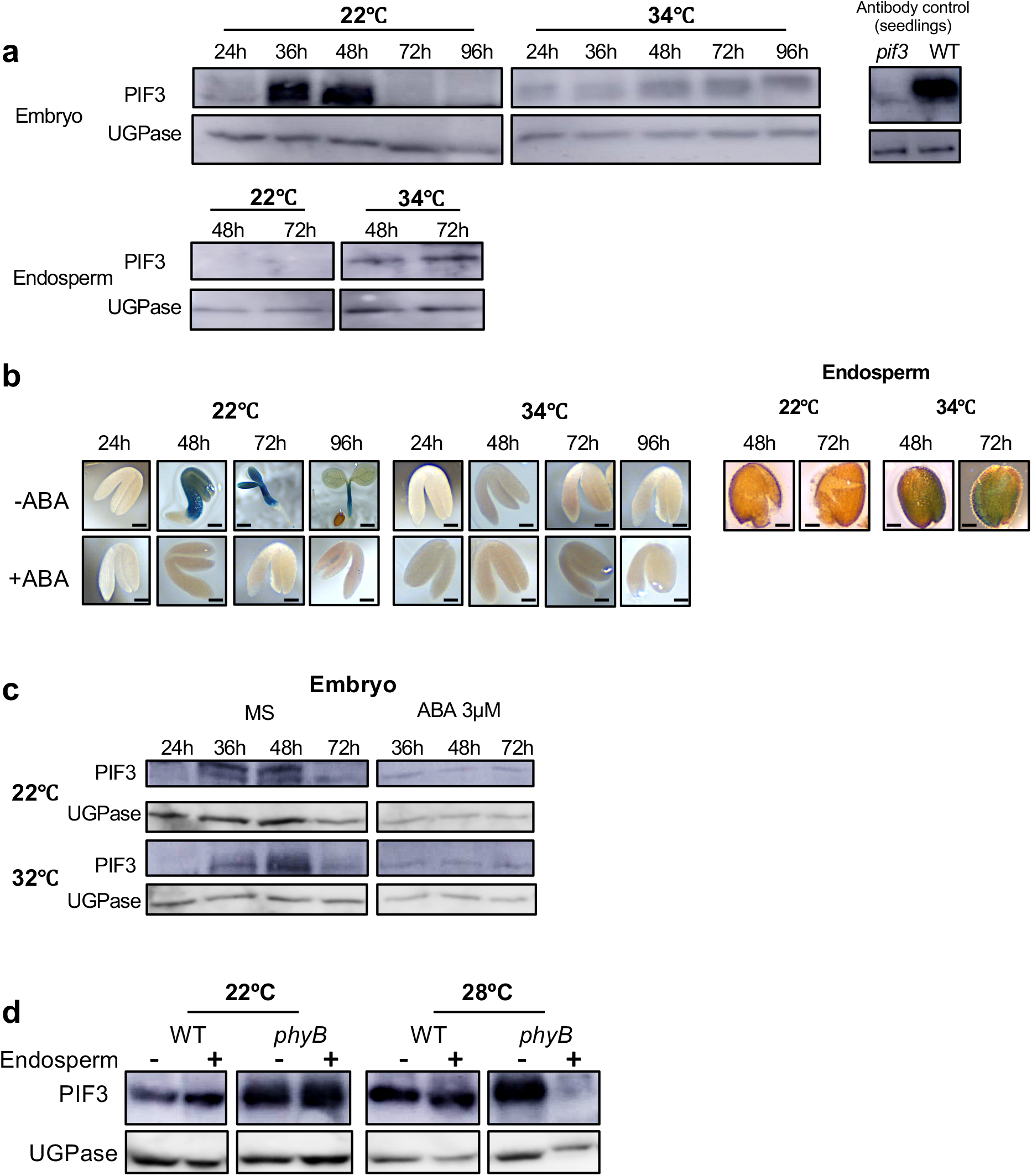
PIF3 activity in the endosperm represses *PIF3* expression in the embryo. a. Protein gel blot analysis of PIF3 levels in WT embryos and endosperms cultivated at 22 and 34°C and dissected at the indicated times upon seed imbibition. PIF3 antibody control: protein extract from WT and *pif3* seedlings grown for 2 days in darkness. UGPase protein levels were used as a loading control. b. Representative pictures of transgenic embryos and endosperms carrying a *PIF3* promoter fused to GUS dissected at the indicated times upon seed imbibition and stained with X-Gluc. Seeds were cultivated at 22°C or 34°C in absence (-ABA) or presence (+ABA) of 3µM ABA. Bar: 0.1 mm for all pictures except MS 48h (bar: 0.25 mm), 72h (bar 0.5 mm) and 96 h (bar 0.75 mm). c. PIF3 protein levels in WT embryos cultivated at 22°C or 32°C in absence (MS) or presence of 3µM ABA and dissected at the indicated times upon seed imbibition. UGPase protein levels were used as a loading control. d. Protein gel blot analysis of PIF3 levels in WT and *phyB* embryos that were either dissected 4 hours after seeds imbibition and cultivated for 3 days (-) or dissected after seeds were cultivated for 3 days (+) at 22°C or 28°C. UGPase protein levels were used as a loading control.

In absence of endosperm, embryos cultured at 22°C and 34°C similarly accumulated transiently PIF3 (Fig. 5c). This suggested that ABA released by the endosperm represses PIF3 accumulation in the embryo. Consistent with this notion, embryos cultured in presence of exogenous ABA maintained low PIF3 levels over time at both 22°C and 34°C (Fig. 5c). Furthermore, exogenous ABA also strongly reduced GUS activity in prom*PIF3::GUS* transgenic embryos (Fig. 5b). In addition, 48h upon imbibition at 28°C, a temperature where *phyB* seeds are thermoinhibited, phyB embryos surrounded by the endosperm accumulated markedly lower PIF3 levels than embryos deprived of their endosperm (Fig. 5d). No such differences in PIF3 accumulation were observed at 22°C (Fig. 5d). Altogether, these results show that under high temperatures endospermic ABA inhibits the accumulation of *PIF3* mRNA and PIF3 protein in the embryo.

### PIF3 regulates largely different sets of genes in the endosperm and embryo

We compared the transcriptome of WT and *pif3* endosperm from seeds imbibed for 48h at 30°C, i.e. at a time when WT and *pif3* seeds are developmentally indistinguishable as no *pif3* germination as yet taken place (Supplementary Figure 3a).We also compared the transcriptome of WT and *pif3* embryos cultured for 48h at 30°C after removing their endosperm 4h upon seed imbibition, which is a time when WT and *pif3* hypocotyl length was comparable (Supplementary Figure 3a).

We found that 1168 and 1186 genes were up- and down-regulated, respectively, in *pif3* endosperm relative to WT endosperm while only 158 and 107 were down- and up-regulated, respectively, in the *pif3* embryo relative to the WT embryo (Supplementary Figure 6b and Supplementary Table 1). Only 16 genes were regulated by PIF3 in both embryo and endosperm (Figure 6b). These data indicate that PIF3 regulates largely different sets of genes in the endosperm and embryo in seeds exposed to high temperatures and that PIF3 has a comparatively much higher impact on endosperm gene expression under higher temperatures, consistent with its role to promote thermoinhibition. A gene ontology (GO) enrichment analysis revealed that the most obvious categories for down-regulated endospermic genes in *pif3* relative to WT are associated with RNA metabolism (including rRNA, snoRNA and ncRNA metabolism) and ribosome biogenesis, whereas those for up-regulated endospermic genes are associated with cell wall modifications (Supplementary Table 1). Among endospermic ABA synthesis genes, we found that 7 of them had their expression significantly deregulated between WT and *pif3* endosperms. The highest differences were observed in genes involved in ABA inactivation (Supplementary Table 1). In particular, we found that the expression of *CYP707A1*, encoding a key catabolic ABA 8’-hydroxylase in seeds, was 6-fold upregulated in *pif3* endosperm at 30°C relative to WT endosperm and, to a lesser extent, also that of *CYP707A3* (2.5 fold)(Supplementary Table 1) ^29^.

**Figure 6.**
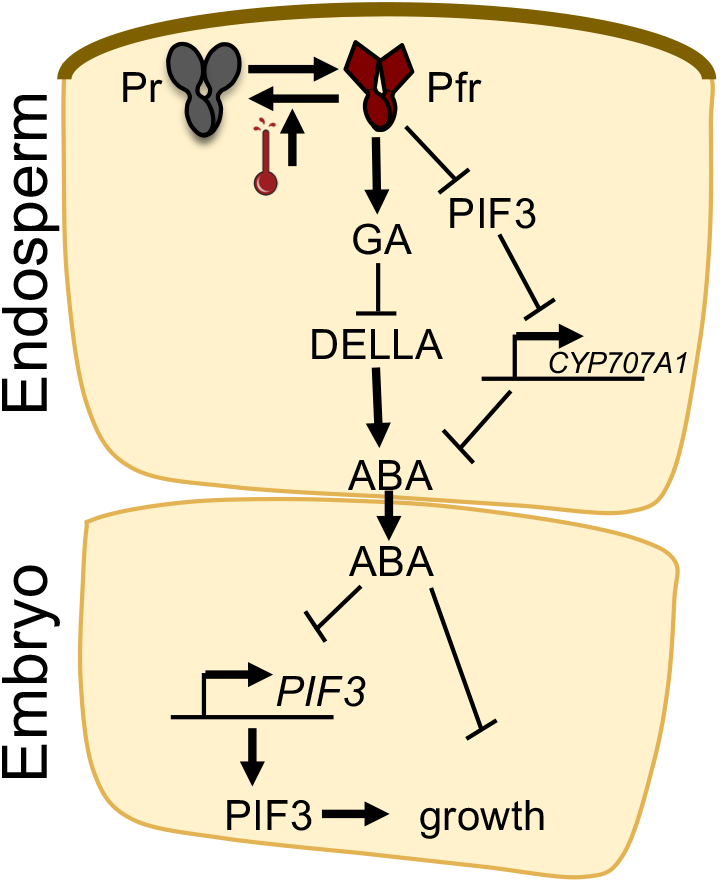
A model for seed thermoinhibition in *Arabidopsis thaliana*. High temperatures accelerate thermal reversion of phyB, which leads to lower levels of its Pfr signaling active form. This activates two parallel pathways promoting endospermic ABA synthesis and release towards the embryo to promote thermoinhibition by repressing growth, which includes repressing the expression of embryonic *PIF3* whose product promotes growth. In one pathway, GA synthesis is repressed, a process that may involve PIF factors (not shown in the model), which leads to stabilization of DELLA factors that promote ABA synthesis. In another pathway, stabilization of PIFs, and particularly PIF3, leads to increased ABA synthesis, which may involve in particular PIF3-dependent repression of *CYP707A1* expression.

## Discussion

Here we showed that the endosperm is essential for seed thermoinhibition to take place in *Arabidopsis thaliana*. Our results are consistent with the model that high temperatures inhibit phyB signaling by accelerating the reversion of the Pfr active form of phyB into its Pr inactive form, as previously shown in seedlings ^12, 13^. Our results indicate that low endospermic phyB signaling promotes endospermic ABA synthesis and release via two parallel signaling branches: one involving the DELLA factors and another involving the PIFs factors and particularly PIF3 (Figure 6). This model does not exclude that each branch could also regulate the activity of the other. Indeed, previous work using whole seeds showed that PIF1 represses and promotes GA synthesis and *DELLA* gene expression, respectively ^30, 31^. Thus, PIF1 or other PIFs could similarly influence GA signaling responses in the endosperm under high temperatures. Furthermore, DELLA and PIFs factors are known to interact with each other, modulating each other’s stability and activity ^10, 32^.

Previous work showed that DELLA factors promote ABA synthesis, which we further confirmed here in the context of seed thermoinhibition. However, we provided evidence that PIFs also promote ABA synthesis and thermoinhibition independently of DELLA factors. In the case of PIF3, this could involve PIF3-mediated repression of *CYP707A1* expression. Furthermore, PIF1 was shown to promote ABA synthesis by promoting expression of the ABA biosynthesis genes *NINE-CIS-EPOXYCAROTENOID DIOXYGENASE6* (*NCED6*) and *NCED9* and repressing that of *CYP707A2*, encoding another important catabolic ABA 8’-hydroxylase in seeds ^29,30^.

Previous work has implicated PIF4 in several processes regulated by temperature such as promoting hypocotyl and petiole growth in seedlings or flowering in response to high temperatures ^8, 9^. Concerning seed thermoinhibition, we found that PIF4 plays a lesser role to promote seed thermoinhibition relative to PIF3 since its contribution is observed only in a *pif3* mutant background (Figure 4a). We observed that endospermic expression of *PIF3* and *PIF4* is strongly induced by high temperatures, however, *PIF3* mRNA accumulation reaches markedly higher levels than that of *PIF4*, being 6-fold higher at 30°C and 2.5-fold higher at 34°C (Supplementary Figure 4). Thus, the importance of endospermic PIF3 relative to other PIFs could be explained, at least in part, by a higher PIF3 protein accumulation relative to other PIFs as a result of higher *PIF3* mRNA accumulation in response to high temperatures.

Under high temperatures, we found that *pif3* endosperm has lower expression in genes associated with RNA metabolism and ribosome biogenesis whereas it has higher expression in genes associated with cell wall modifications (Supplementary Table 1). These observations might be consistent with the notion that *pif3* endospermic cells are in a distinct metabolic state relative to that of WT endospermic cells. Indeed, in *pif3* seeds, which lack thermoinhibition, the *pif3* endosperm does not repress germination and is therefore bound to degenerate as it is abandoned by the seedling after germination. This could explain the drop in RNA metabolism gene expression. On the other hand, the increase in cell wall modification gene expression might reflect that the *pif3* endosperm cells are bound to detach from each other to allow the embryonic radicle to emerge from the seed (germination).

Seed thermoinhibition can only be observed in non-dormant seeds since dormant seeds will not germinate under normal favorable conditions, i.e. imbibition at 22°C in presence of white light (normal conditions). In the Col-0 accession, dormancy is lost after a couple of weeks of dry after-ripening, i.e. two-week old WT Col-0 seeds will germinate at 22°C. Interestingly, seed thermoinhibition is strongest in younger WT seed batches. For example, a two-week-old WT seed batch will be fully thermoinhibited at 28°C, unlike a two-month-old WT seed batch. If thermoinhibition results from higher temperatures lowering phyB signaling in the endosperm and if thermoinhibition at a given temperature becomes weaker with older seeds, then the model predicts that phyB-mediated germination ought to gain strength as seed age. This notion is indeed supported by experiments performed by De Giorgi et al. using WT seeds of various seed ages ^33^. Indeed, in two-week-old seeds able to germinate under normal conditions, a pulse of red light early upon seed imbibition followed by incubation in darkness, which normally leads to phyB-mediated germination, is insufficient to promote germination ^33, 34^. In two-month-old seeds, the same treatment only partially triggers germination and full germination is only observed in 6-month-old WT seeds ^33^.

If thermoinhibition is the result of lower phyB signaling, then one would expect *phyB* mutant seeds to never germinate even under normal conditions. This is indeed the case to some degree: two-week old *phyB* mutants are unable to germinate under normal conditions (22°C), unlike WT seeds. Full *phyB* germination under normal conditions is only observed in two-month-old *phyB* seeds. In this study, we therefore systematically used seed batches that are at least two-month-old so that WT and *phyB* germination is comparable under normal germination conditions. Two-month-old *phyB* seeds could germinate under normal conditions but not under darkness (Supplementary Figure 5a). This suggests that the other phytochromes promote seed germination at 22°C under normal conditions and high temperatures would also accelerate their reversion to their inactive state in *phyB* mutants exposed to 28°C thus leading to *phyB* mutant thermoinhibition. Consistent with this view, two-month-old *phyAphyB* and *phyBphyCphyDphyE* seeds were unable to germinate at 22°C (Supplementary Figure 5b) whereas three-year-old *phyAphyB* seeds germinated like WT seeds and *phyBphyCphyDphyE* seeds did not germinate well (Supplementary Figure 5c). Furthermore, three-year-old *phyAphyB* and *phyBphyCphyDphyE* seeds were thermoinhibited at 28°C (Supplementary Figure 5c). However, the contribution of the other phytochromes for seed thermoinhibition could only be revealed in absence of *phyB* as *phyAphyCphyDphyE* seeds behaved similar to WT seeds (Supplementary Figure 5b and c). Altogether, these observations indicate temperature-driven phyB inactivation plays an essential role to promote thermoinhibition, which is reinforced by the inactivation of the remaining phytochromes.

Previous work provided indirect evidence that endospermic PIF1 promotes ABA synthesis and release from the endosperm to repress germination in response to farred light ^14^. Here, we showed that PIF3 indeed promotes ABA synthesis and release under higher temperatures to promote thermoinhibition. Hence, the work presented here helps understanding a paradox concerning how PIFs regulate early seedling development. Indeed, whereas PIFs promote growth in seedlings, some PIFs, such as PIF1 and PIF3, repress embryonic growth in seeds. This paradox is partly resolved if one considers that the role of PIFs to repress germination reflects their activity to promote endospermic ABA synthesis and release in the endosperm rather than their activity to promote growth in the embryo. Interestingly, PIF3-mediated endospermic ABA release maintains embryonic *PIF3* expression low thus inhibiting embryonic PIF3-mediated growth. Hence, low endospermic phyB signaling hijacks low embryonic phyB signaling, which would stabilize embryonic PIF3, by reducing embryonic *PIF3* expression and thus preventing embryonic PIF3 accumulation. Our work here consolidates the importance of phyB signaling in the endosperm to control the onset of the embryo-to-seedling transition and further reveals the importance of PIFs as essential phyB signaling components also in the endosperm. phyB signaling was mainly studied and characterized in seedlings where it regulates numerous developmental processes ^10, 11^. The study of endospermic phyB signaling and the identification of its specificities will enable to understand how evolution has tinkered with phyB-mediated signaling pathways to implement germination control mechanisms in angiosperms, producing endosperm-bearing seeds, while retaining adaptive phyB signaling developmental responses in seedlings.

## Acknowledgments

We thank Pablo Cerdan for providing *phyACDE, phyBCDE* and *phyAB* seeds, Elena Monte for *pif3* and WT/*promPIF3:GUS* seeds, Ferenc Nagy for *phyB* mutant seeds complemented with PHYB that has a WT or Ser86Asp or Gly564Glu sequences, Akira Nagatani for providing phyB antibody. We would like to thank Mylene Docquier and members of the Genomics Platform of the Institute of Genetics and Genomics (iGE3) at the University of Geneva for help with RNAseq experiments. We specially thank Mayumi Iwasaki for discussions, help with bioinformatics and critical reading of the manuscript.

## Material and Methods

### Plant material

The *Arabidopsis* mutant and transgenic seeds used in this study (Col-0 background) were described previously: *aba1-6* ^35^, *della* (*rgl2-Sk54 rga-28 gai-t6 rgl1-Sk62 rgl3-3*) ^36^, *phyA-211* ^37^, *phyB-9* ^38^, *phyB-9/phyC-2/phyD-201/phyE-201* and *phyA-211/phyC-2/phyD-201/phyE-201* ^39^, *pif3-1 and pif3-3* ^27^, *pif3-3/phyB-9, pif1-1/pif3-3* and *pif1-1/pif3-3/pif5-3* ^40^, *pif1-1* ^15^, *pif4-2* and *pif7-1* ^40^, *pif5-3* ^41^ and *pif6-1* (SALK_090239C). *phyB-9*/*35S:phyB-YFP* and *phyB-9*/*35S:phyB*^*S86D*^-YFP ^19^, WT/35S:*phyB-GFP* and WT/35S:phyB^G564E^-GFP ^20^, WT/*promPIF3:GUS* ^42^. The mutant *phyB/della* and *pif3/della* mutants were created using CRISPR/Cas9 editing of the *della* mutant. The *PHYB* mutant allele has an A insertion at position 603 leading to a frameshift and the *PIF3* mutant allele has C insertion at position 1847 leading to a frameshift.

### Plant growth conditions and germination assays

The *Arabidopsis* mutant and transgenic seeds were harvested on the same day from plants grown under the same conditions and after-ripened at room temperature for the same time period. For the germination assays, seeds were surface sterilized and sown on MS medium (Sigma) containing 0.8% (w/v) agar. R and FR light treatment was performed as previously described ^43^. The Seed Coat Bedding Assay was performed as previously described using 30 endosperms for 1 embryo or 100 endosperms for 4 embryos (Figure 2c and d) ^14^.

### Generation of *phyB*/*della* and *pif3*/*della* mutants using CRISPR/Cas9 genome editing

For the generation of CRISPR/Cas9 mutant alleles of *PHYB* and *PIF3* in the *della* mutant, the method described by Tsutsui and Higashiyama was used ^44^. For targeting the 5’ end of *PHYB*, the primers 5’-attgatctgttgctcaggtacag -3’ and 5’-aaacctgtacctgagcaacagat -3’ were annealed and cloned into the pKIR1.1 plasmid. For targeting the 5’ end of *PIF3*, the primers 5’-attgagcaatatccatcaaggga -3’ and 5’-aaactcccttgatggatattgct-3’ were used.

### RNA extraction and RT-qPCR

Total RNA was extracted from 200 endosperms or embryos as previously described ^45^. Total RNA was treated with RQ1 RNase-Free DNase (Promega) and reverse-transcribed using ImProm II reverse transcriptase (Promega) and oligo(dT)15 primer. Quantitative RT-PCR was performed using Quant Studio 5 real-time PCR system (Applied Biosystems) and GoTaq qPCR Master Mix (Promega). Relative transcript level was calculated using the comparative Δ Ct method and normalized to the *PP2A* gene transcript levels. The primers to amplify *CYP707A1* (At4g19230) are 5’-tcatctcaccaccaagta-3’ and 5’-aaggcaattctgtcattcta -3’ ^46^; *PIF3* (At1g09530) 5’-gacggcgtgataggatcaac-3’ and 5’-catcgaagctttgtccacct-3’; *PIF4* (At2g43010) 5’-aagtcgaaccaacgatcagg-3’ and 5’-ttgcaaagccttcattctctc-3’ ^47^; *PP2A* (At1g69960) 5’-ggaccggagccaactagga-3’ and 5’-gctatccgaacttctgcctcatt-3’ ^14^.

### Histochemical GUS staining

Histochemical GUS staining assay was performed using a substrate buffer: 100 mM sodium phosphate (pH 7.0), 5 mM potassium ferricyanide, 5 mM potassium ferrocyanide, 1mM EDTA, 1% Triton-X, 1 mg/mL X-Gluc for embryo and 5 mg/mL for endosperm samples. Samples were incubated at 37ºC for 16 hours.

### Antibody production and protein gel blot

PIF3 recombinant proteins were prepared using PIF3-his DNA (pET28C, Novagen) provided by Ferenc Nagy and induced and purified using commercial kit (Amersham). Polyclonal anti-PIF3 was obtained from rabbits immunized with PIF3 recombinant protein. PIF3 antibodies were further affinity-purified using PIF3 recombinant protein immobilized on nitrocellulose filters as described ^6^. Endosperms (200) and embryos (20) were homogenized in presence of protein extraction buffer (50 mM Tris-HCl pH 6.8, 2% SDS, 10% glycerol, 100 mM DTT, 0.01% bromophenol blue) and protein extract corresponding to 150 endosperms or 15 embryos was loaded per lane. Proteins were separated on 10% SDS-PAGE gel and transferred to a PVDF membrane (Amersham). Membranes were incubated in 5% milk in Tris-buffered saline (TBS) containing PIF3 antibody (5:1,000 dilution), RGL2 antibody (4:1,000 dilution) ^6^ or ABI5 antibody (5:10,000 dilution) ^17^ for 16 hours at 4ºC. Anti-UGPase antibody (Agrisera) was used at 1:10,000 dilution and anti-PHYB antibody and used at 1:1,000 dilution for 16 hours at 4ºC. Anti-rabbit (for PIF3, RGL2, ABI5 and UGPase) or antimouse (for PHYB) IgG HRP-linked whole antibody (GE healthcare) in a 1:10,000 dilution was used as secondary antibody for 2 hours at RT. Membranes were washed with TBS+ 0.05% Tween 3 times for 10 minutes after the first and second antibody incubation and the immune complexes were detected using the ECL kit (Amersham).

### RNA-seq and gene expression profile analysis

Total RNA was isolated from 200 endosperms dissected 40 hours after seeds incubation at 30ºC and from 200 embryos dissected 4 hours after seeds imbibition and growing for 40 hours at 30ºC. Total RNA was isolated as previously described ^45^. cDNA libraries were prepared from 200ng of total RNA using a TruSeq mRNA Library Prep Kit (Illumina). cDNA libraries were normalized and pooled then sequenced using HiSeq 2500 (Illumina) with single-end 50 bp reads. Transcript assembly and normalization was performed with the Cufflinks program and gene expression levels were calculated in FPKM (Fragments Per Kilobase of exon per Milion mapped fragments) units. Differential gene expression analysis was performed by Cuffdiff, a part of the Cufflinks package ^48, 49, 50^.

### ABA measurement

ABA measurements were performed using a protocol adapted from Glauser et al. ^51^. For the extraction of supernatants, 10 ul of ABA-d6 (100 ng/mL in water) were added to the collected supernatant and the mixture was lyophilized overnight. The dry residue was resuspended in 100 ul of methanol 35%., vortexed and sonicated for 1 min each, and the suspension was transferred in a 0.2 mL PCR tube and centrifuged at 14’000 g for 2 min. The supernatant was collected and placed in an HPLC vial fitted with a conical insert for analysis. The extraction of plant tissues was done as follows. The tissue was lyophilized overnight in a 2.0 mL Eppendorf tube. Then the tube was frozen in liquid nitrogen and ground into powder using two 3mm stainless steel beads in a mixer mill (30 Hz, 15 s). 990 ul of ethylacetate:formic acid (99.5:0.5, v/v) and 10 ul of ABA-d6 (100 ng/mL in water) were added and the tube was shaken for 3 min at 30 Hz. After centrifugation at 14’000 g for 3 min, the supernatant was transferred to a new 2.0 mL tube, and the pellet from the first tube was re-extracted using 0.5 mL of ethylacetate:formic acid (99.5:0.5, v/v). Both supernatants were combined and evaporated to dryness at 35°C in a centrifugal evaporator. The residue was resuspended in 100 ul of methanol 35%, vortexed and sonicated for 1 min each, and the suspension was transferred in a 0.2 mL PCR tube and centrifuged at 14’000 g for 2 min. The supernatant was collected and placed in an HPLC vial fitted with a conical insert for analysis.

The analysis of ABA was done by HPLC-MS/MS using a QTRAP 6500+ (Sciex) connected to an Acquity UPLC (Waters). 3.7 ul of extract was injected onto an Acquity UPLC BEH C18 column (50×2.1mm, 1.7 um particle size, Waters). A gradient of phase A - H2O:formic acid 0.05% and phase B - acetonitrile:formic acid 0.05% from 5-65% B in 6.5 min was applied. The flow rate was set to 0.4 mL/min and the column temperature to 35 °C. The mass spectrometer was operated in electrospray negative ionization using the multiple reaction monitoring (MRM) mode. Transitions for ABA and ABA-d6 were 263/153 and 269/159, respectively. A five-point calibration curve (0.02, 0.1, 0.5, 5 and 20 ng/mL, all containing ABA-d6 at 10 ng/mL) was used for quantification. Linear regressions weighted by 1/x were applied. Analyst v.1.7.1 was used to control the instrument and for data processing. We ran blank samples and 0.02 ng/mL ABA standards over the run of the batches as quality controls.

## References

1. Iwasaki M, Penfield S, Lopez-Molina L. Parental and Environmental Control of Seed Dormancy in Arabidopsis thaliana. Annu Rev Plant Biol, (2022).

2. Toh S, et al. High temperature-induced abscisic acid biosynthesis and its role in the inhibition of gibberellin action in Arabidopsis seeds. Plant Physiol 146, 1368–1385 (2008).

3. Huo H, Dahal P, Kunusoth K, McCallum CM, Bradford KJ. Expression of 9-cis-EPOXYCAROTENOID DIOXYGENASE4 is essential for thermoinhibition of lettuce seed germination but not for seed development or stress tolerance. Plant Cell 25, 884–900 (2013).

4. Tamura N, et al. Isolation and characterization of high temperature-resistant germination mutants of Arabidopsis thaliana. Plant Cell Physiol 47, 1081–1094 (2006).

5. Yoong FY, et al. Genetic Variation for Thermotolerance in Lettuce Seed Germination Is Associated with Temperature-Sensitive Regulation of ETHYLENE RESPONSE FACTOR1 (ERF1). Plant Physiol 170, 472–488 (2016).

6. Piskurewicz U, Jikumaru Y, Kinoshita N, Nambara E, Kamiya Y, Lopez-Molina L. The gibberellic acid signaling repressor RGL2 inhibits Arabidopsis seed germination by stimulating abscisic acid synthesis and ABI5 activity. Plant Cell 20, 2729–2745 (2008).

7. Piskurewicz U, Tureckova V, Lacombe E, Lopez-Molina L. Far-red light inhibits germination through DELLA-dependent stimulation of ABA synthesis and ABI3 activity. EMBO J 28, 2259–2271 (2009).

8. Koini MA, et al. High temperature-mediated adaptations in plant architecture require the bHLH transcription factor PIF4. Curr Biol 19, 408–413 (2009).

9. Kumar SV, et al. Transcription factor PIF4 controls the thermosensory activation of flowering. Nature 484, 242–245 (2012).

10. Cheng MC, Kathare PK, Paik I, Huq E. Phytochrome Signaling Networks. Annu Rev Plant Biol 72, 217–244 (2021).

11. Legris M, Ince YC, Fankhauser C. Molecular mechanisms underlying phytochrome-controlled morphogenesis in plants. Nature communications 10, 5219 (2019).

12. Jung JH, et al. Phytochromes function as thermosensors in Arabidopsis. Science 354, 886–889 (2016).

13. Legris M, et al. Phytochrome B integrates light and temperature signals in Arabidopsis. Science 354, 897–900 (2016).

14. Lee KP, et al. Spatially and genetically distinct control of seed germination by phytochromes A and B. Genes Dev 26, 1984–1996 (2012).

15. Oh E, Kim J, Park E, Kim J-I, Kang C, Choi G. PIL5, a Phytochrome-Interacting Basic Helix-Loop-Helix Protein, Is a Key Negative Regulator of Seed Germination in Arabidopsis thaliana. Plant Cell 16, 3045–3058 (2004).

16. Lee KP, Piskurewicz U, Tureckova V, Strnad M, Lopez-Molina L. A seed coat bedding assay shows that RGL2-dependent release of abscisic acid by the endosperm controls embryo growth in Arabidopsis dormant seeds. Proc Natl Acad Sci U S A 107, 19108–19113 (2010).

17. Lopez-Molina L, Mongrand S, Chua NH. A postgermination developmental arrest checkpoint is mediated by abscisic acid and requires the ABI5 transcription factor in Arabidopsis. Proc Natl Acad Sci U S A 98, 4782–4787 (2001).

18. Heschel MS, Selby J, Butler C, Whitelam GC, Sharrock RA, Donohue K. A new role for phytochromes in temperature-dependent germination. New Phytol 174, 735–741 (2007).

19. Medzihradszky M, et al. Phosphorylation of phytochrome B inhibits light-induced signaling via accelerated dark reversion in Arabidopsis. Plant Cell 25, 535–544 (2013).

20. Adam E, et al. Altered dark- and photoconversion of phytochrome B mediate extreme light sensitivity and loss of photoreversibility of the phyB-401 mutant. PloS one 6, e27250 (2011).

21. Kretsch T, Poppe C, Schafer E. A new type of mutation in the plant photoreceptor phytochrome B causes loss of photoreversibility and an extremely enhanced light sensitivity. Plant J 22, 177–186 (2000).

22. Yamaguchi S, Smith MW, Brown RG, Kamiya Y, Sun T. Phytochrome regulation and differential expression of gibberellin 3beta-hydroxylase genes in germinating Arabidopsis seeds. Plant Cell 10, 2115–2126 (1998).

23. Griffiths J, et al. Genetic characterization and functional analysis of the GID1 gibberellin receptors in Arabidopsis. Plant Cell 18, 3399–3414 (2006).

24. McGinnis KM, et al. The Arabidopsis SLEEPY1 gene encodes a putative F-box subunit of an SCF E3 ubiquitin ligase. Plant Cell 15, 1120–1130 (2003).

25. Nakajima M, et al. Identification and characterization of Arabidopsis gibberellin receptors. Plant J 46, 880–889 (2006).

26. Willige BC, et al. The DELLA domain of GA INSENSITIVE mediates the interaction with the GA INSENSITIVE DWARF1A gibberellin receptor of Arabidopsis. Plant Cell 19, 1209–1220 (2007).

27. Monte E, et al. The phytochrome-interacting transcription factor, PIF3, acts early, selectively, and positively in light-induced chloroplast development. Proc Natl Acad Sci U S A 101, 16091–16098 (2004).

28. Stavang JA, et al. Hormonal regulation of temperature-induced growth in Arabidopsis. Plant J 60, 589–601 (2009).

29. Okamoto M, et al. CYP707A1 and CYP707A2, which encode abscisic acid 8’-hydroxylases, are indispensable for proper control of seed dormancy and germination in Arabidopsis. Plant Physiol 141, 97–107 (2006).

30. Oh E, et al. PIL5, a phytochrome-interacting bHLH protein, regulates gibberellin responsiveness by binding directly to the GAI and RGA promoters in Arabidopsis seeds. Plant Cell 19, 1192–1208 (2007).

31. Paik I, Kathare PK, Kim JI, Huq E. Expanding Roles of PIFs in Signal Integration from Multiple Processes. Molecular plant 10, 1035–1046 (2017).

32. Feng S, et al. Coordinated regulation of Arabidopsis thaliana development by light and gibberellins. Nature 451, 475–479 (2008).

33. De Giorgi J, et al. An Endosperm-Associated Cuticle Is Required for Arabidopsis Seed Viability, Dormancy and Early Control of Germination. PLoS genetics 11, e1005708 (2015).

34. Shinomura T, Nagatani A, Chory J, Furuya M. The Induction of Seed Germination in Arabidopsis thaliana Is Regulated Principally by Phytochrome B and Secondarily by Phytochrome A. Plant Physiol 104, 363–371 (1994).

35. Barrero JM, et al. A mutational analysis of the ABA1 gene of Arabidopsis thaliana highlights the involvement of ABA in vegetative development. J Exp Bot 56, 2071–2083 (2005).

36. Park J, Nguyen KT, Park E, Jeon JS, Choi G. DELLA proteins and their interacting RING Finger proteins repress gibberellin responses by binding to the promoters of a subset of gibberellin-responsive genes in Arabidopsis. Plant Cell 25, 927–943 (2013).

37. Reed JW, Nagatani A, Elich TD, Fagan M, Chory J. Phytochrome A and Phytochrome B Have Overlapping but Distinct Functions in Arabidopsis Development. Plant Physiol 104, 1139–1149 (1994).

38. Reed JW, Nagpal P, Poole DS, Furuya M, Chory J. Mutations in the gene for the red/far-red light receptor phytochrome B alter cell elongation and physiological responses throughout Arabidopsis development. Plant Cell 5, 147–157 (1993).

39. Sanchez-Lamas M, Lorenzo CD, Cerdan PD. Bottom-up Assembly of the Phytochrome Network. PLoS genetics 12, e1006413 (2016).

40. Leivar P, et al. Multiple phytochrome-interacting bHLH transcription factors repress premature seedling photomorphogenesis in darkness. Curr Biol 18, 1815–1823 (2008).

41. Khanna R, et al. The basic helix-loop-helix transcription factor PIF5 acts on ethylene biosynthesis and phytochrome signaling by distinct mechanisms. Plant Cell 19, 3915–3929 (2007).

42. Zhang Y, et al. A quartet of PIF bHLH factors provides a transcriptionally centered signaling hub that regulates seedling morphogenesis through differential expression-patterning of shared target genes in Arabidopsis. PLoS genetics 9, e1003244 (2013).

43. Kim W, Zeljkovic SC, Piskurewicz U, Megies C, Tarkowski P, Lopez-Molina L. polyamine uptake transporter 2 (put2) and decaying seeds enhance phyA-mediated germination by overcoming PIF1 repression of germination. PLoS genetics 15, e1008292 (2019).

44. Tsutsui H, Higashiyama T. pKAMA-ITACHI Vectors for Highly Efficient CRISPR/Cas9-Mediated Gene Knockout in Arabidopsis thaliana. Plant Cell Physiol 58, 46–56 (2017).

45. Piskurewicz U, Lopez-Molina L. Isolation of genetic material from Arabidopsis seeds. Methods Mol Biol 773, 151–164 (2011).

46. Shu K, et al. ABI4 regulates primary seed dormancy by regulating the biogenesis of abscisic acid and gibberellins in arabidopsis. PLoS genetics 9, e1003577 (2013).

47. Shor E, Paik I, Kangisser S, Green R, Huq E. PHYTOCHROME INTERACTING FACTORS mediate metabolic control of the circadian system in Arabidopsis. New Phytol 215, 217–228 (2017).

48. Blankenberg D, et al. Galaxy: a web-based genome analysis tool for experimentalists. Current protocols in molecular biology / edited by Frederick M Ausubel [et al] Chapter 19, Unit 19 10 11–21 (2010).

49. Giardine B, et al. Galaxy: a platform for interactive large-scale genome analysis. Genome research 15, 1451–1455 (2005).

50. Goecks J, Nekrutenko A, Taylor J, Galaxy T. Galaxy: a comprehensive approach for supporting accessible, reproducible, and transparent computational research in the life sciences. Genome Biol 11, R86 (2010).

51. Glauser G, Vallat A, Balmer D. Hormone profiling. Methods Mol Biol 1062, 597–608 (2014).

