## Supplementary Figures for "The Arabidopsis endosperm is a temperature-sensing tissue that implements seed thermoinhibition through phyB and PIF3"

### Supplementary Figure 1

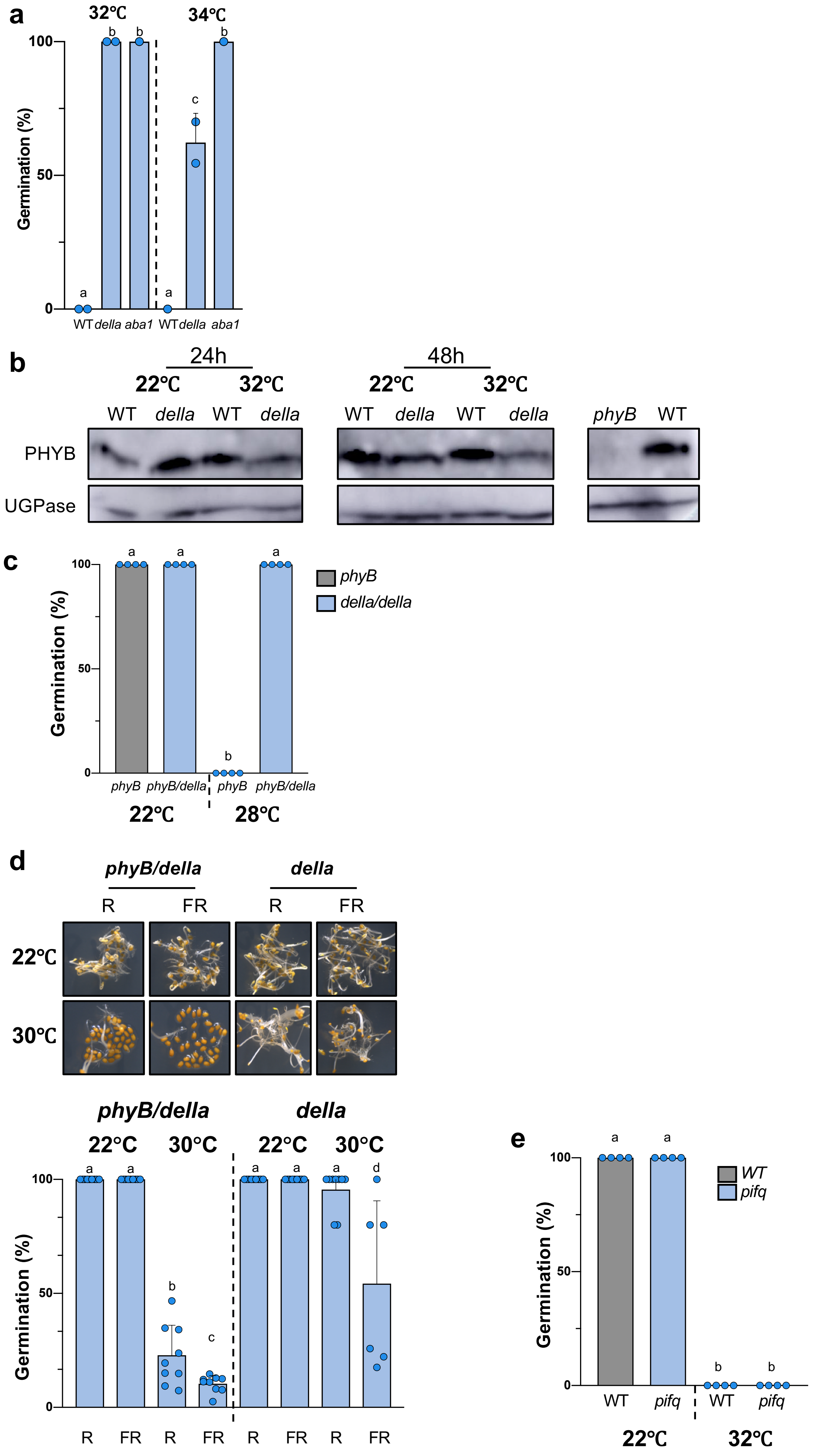

**Supplementary Figure 1**. DELLA factors promote ABA release from endosperm when phyB signaling is low.

a. Average germination percentage of WT, *della* and *aba1* seeds cultured 3 days at 32˚C and 34°C in 3 independent seed batches (n≥50 seeds). Statistical differences assessed by Student`s t-test.

b. PHYB protein levels in WT and *della* seeds cultured at 22˚C or 32°C for 24h and 48 h. UGPase protein levels were used as a loading control. PhyB antibody control: protein extract from WT and *phyB* seedlings grown for 2 days in darkness.

c. Average germination percentage of *phyB* and *phyB*/*della* seeds cultured 3 days at 22˚C and 28˚C in 3 independent seed batches (n≥50 seeds). Statistical differences assessed by Student`s t-test.

d. Representative pictures of *phyB/della* and *della* seeds cultured for 3 days in darkness at 22˚C and 30˚C after receiving a far-red light pulse (FR) or a FR pulse followed by a red light pulse (FR/R) 2h upon seed imbibition. Average germination percentages after 3 days in 5-9 independent seed batches (n≥40 seeds). Statistical treatment and lower-case letters as in Figure 1b.

e. Average germination percentage of WT and *pifq* seeds cultured for 3 days at 22˚C and 32˚C in 3 independent seed batches (n≥50).

### Supplementary Figure 2

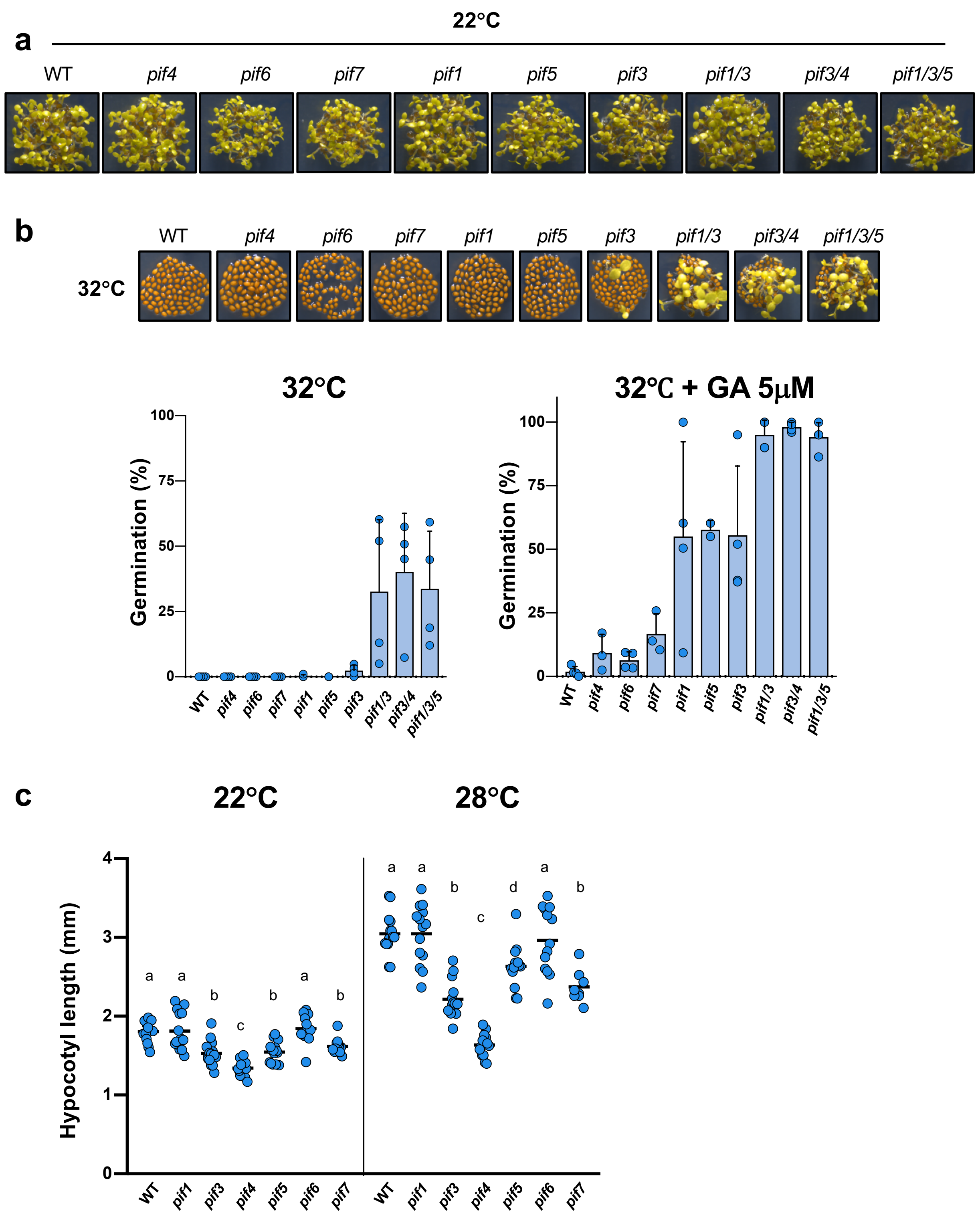


**Supplementary Figure 2**. Endospermic PIF3 promotes seed thermoinhibition by promoting ABA synthesis and release.

a. Representative pictures of WT and *pif* seeds cultured for 3 days at 22°C.

b. Representative pictures of WT and *pif* seeds cultured for 6 days at 32°C. Average germination percentage of WT and *pif* seeds cultured for 6 days at 32°C (left) and for 4 days at 32°C in presence of 5µM GA. 3-4 different seed batches were analyzed (n≥50).

c. Hypocotyl length (mm) in WT and *pifs* seedlings grown according to Koini et al, 2009: seeds were stratified for 3 days and transferred to 22˚C for 4 days and thereupon either kept at 22˚C or transferred at 28˚C for 3 days. n>10 embryos. Statistical treatment and lower-case letters as in Figure 1b.

Supplementary Figure 3

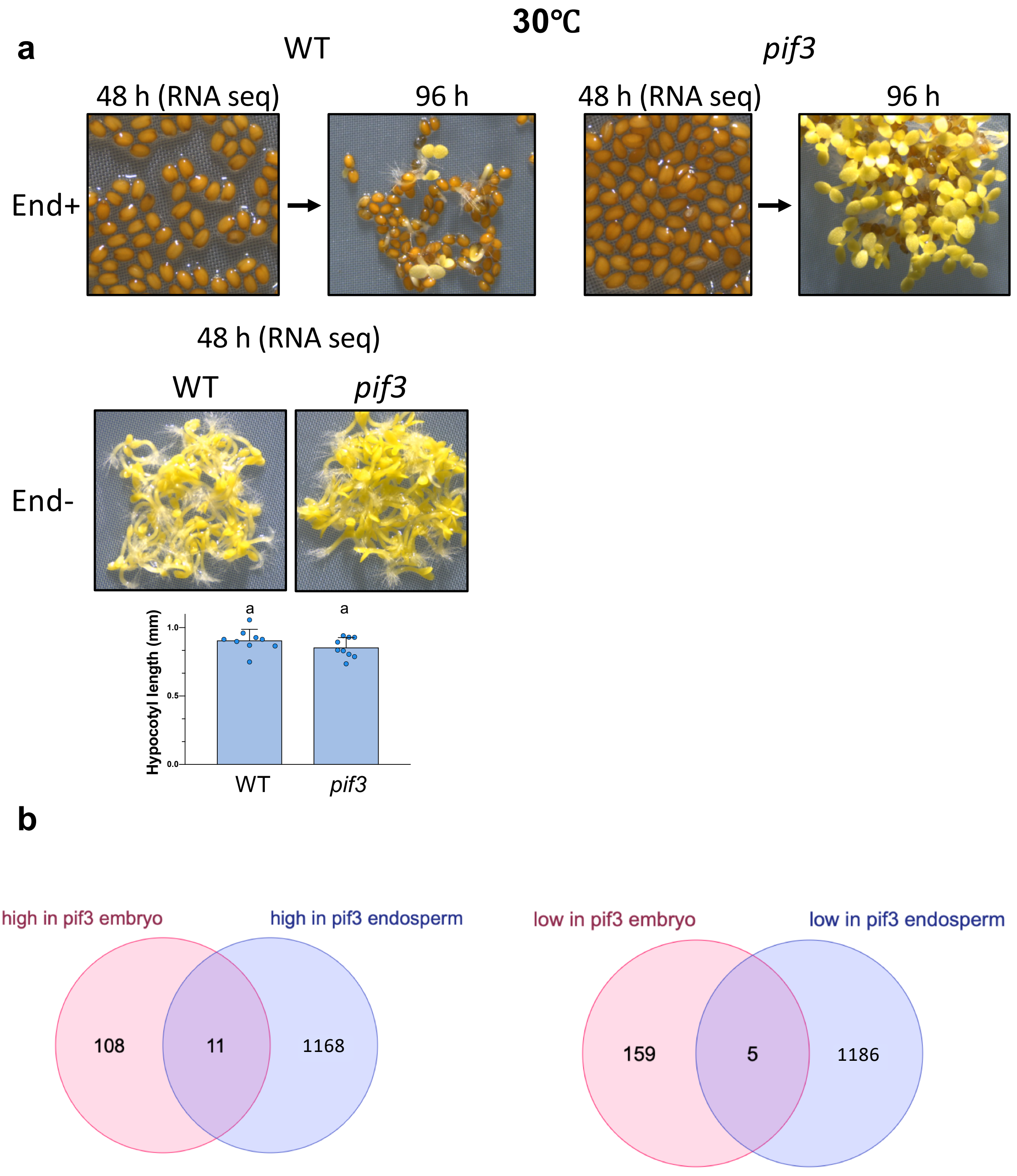


**Supplementary Figure 3**. PIF3 regulates largely different sets of genes in the endosperm and in the embryo.

a. For RNA seq analysis endosperms of WT and *pif3* were collected 48 hours after seeds cultivation at 30°C (End+). Pictures show the time of seeds dissection and endosperms collection (48 h) and the time when *pif3* showed the loss of termoinhibition at 96 hours (upper panel). Pictures below show WT and *pif3* embryos dissected 4 hours after seeds imbibition and incubated for 48 hours at 30°C (End-). Histogram shows hypocotyl length (mm) measured in seedlings used for the RNA seq experiment. n=9 embryos. Statistical treatment and lower-case letters as in Figure 1b.

b. Venn diagrams shows the number of genes that are down-regulated or up-regulated in *pif3* embryo and endosperm relative tp WT embryo and endosperm, respectively.

### Supplementary Figure 4

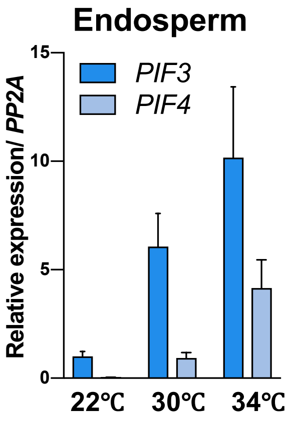


**Supplementary Figure 4.**
Relative *PIF3* and *PIF4* mRNA accumulation in WT endosperm dissected from WT seeds after 3 days of culture at indicated temperatures. *PIF3 and PIF4* expression levels were normalized to those of *PP2A.*

Supplemental figure 5

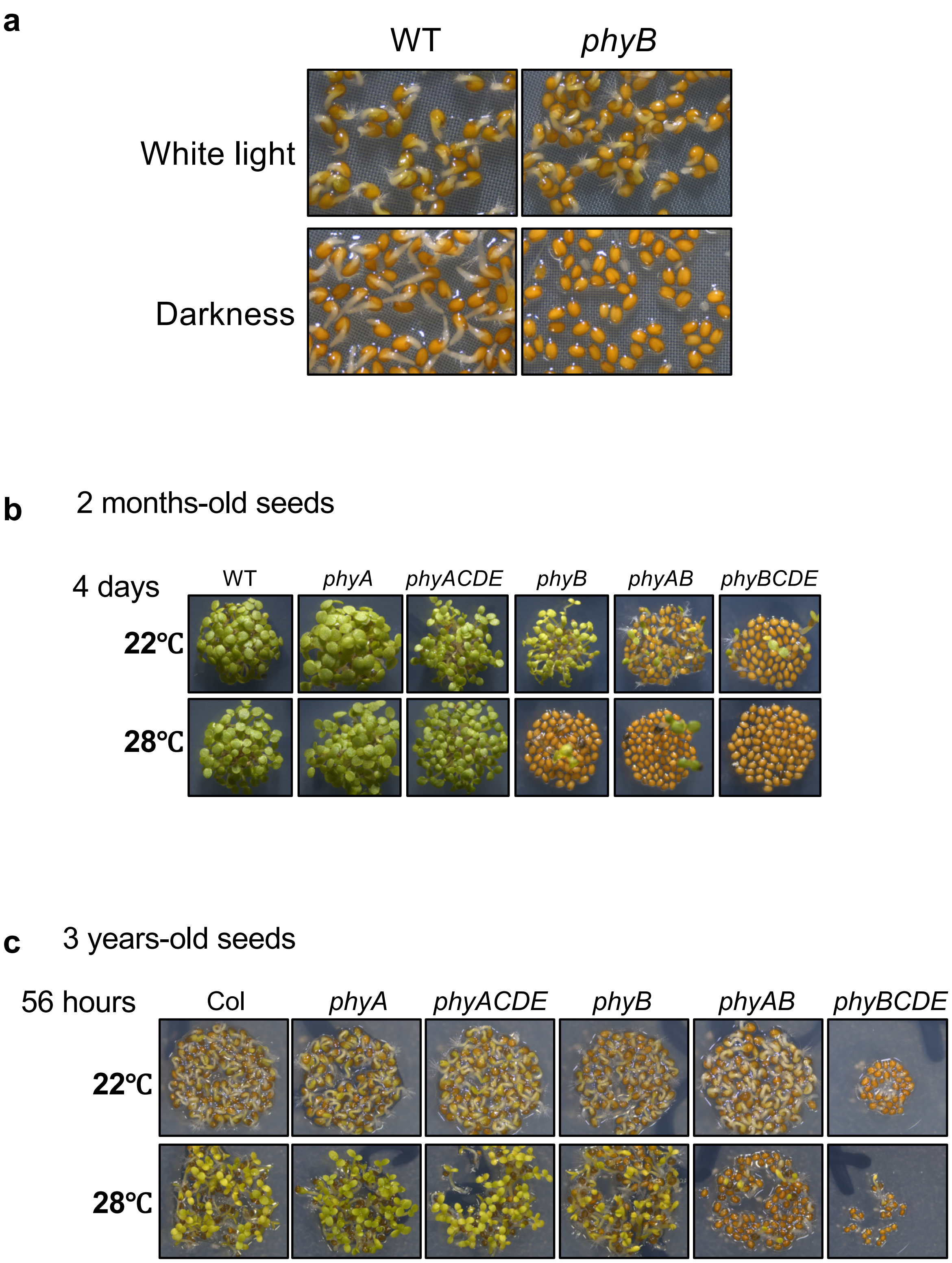


**Supplementary Figure 5**. Phytochrome-mediated germination is enhanced in aging seeds.
a. Representative pictures of WT and *phyB* seeds cultured for 2 days at 22°C in presence or absence of white light.
b. Representative pictures of 2-month old WT, *phyA, phyB, phyAB*, *phyACDE* and *phyBCDE* seeds cultured for 4 days at 22˚C and 28˚C.
c. Same as b. with 3 years-old seeds.
